# Maternal immunity and antibodies to dengue promote Zika virus-induced microcephaly in fetuses

**DOI:** 10.1101/371864

**Authors:** Abhay P. S. Rathore, Wilfried A. A. Saron, Ting Lim, Nusrat Jahan, Ashley L. St John

## Abstract

**Zika virus (ZIKV), a recently emerged flaviviral pathogen, has been linked to microcephaly in neonates^1^. Yet, it is not understood why some fetuses develop severe microcephaly due to maternal ZIKV infection while others do not. The risk for ZIKV-induced microcephaly is greatest during the first trimester of pregnancy in humans^2,3^, yet this alone cannot account for the varied presentation of microcephaly observed. Given the antigenic similarity between ZIKV and closely related dengue virus (DENV)^4^, combined with the substantial immunity to DENV in ZIKV target populations in recent outbreaks, we hypothesized that maternal antibodies against DENV could promote ZIKV-induced microcephaly. Here, using immune-competent mice, we show that maternal to fetal transmission of ZIKV occurs, leading to fetal infection and disproportionate microcephaly. DENV-specific antibodies in pregnant female mice enhance vertical transmission of infection and result in a severe microcephaly like-syndrome during ZIKV infection. Furthermore, fetal infection was promoted by the neonatal Fc receptor (FcRN). Our results identify a novel immune-mediated mechanism of vertical transmission of viral infection and raise caution since ZIKV epidemic regions are also endemic to DENV.**

## Introduction

Zika virus (ZIKV) belongs to the family of *Flaviviridae*, which includes other arboviruses, such as dengue virus (DENV), Japanese encephalitis virus (JEV) and West Nile virus^4^. ZIKV unexpectedly surfaced recently as a major public health concern due to the ongoing and spreading epidemic in South and Central America and the realization that it could cause birth defects and neurological complications^3,5^. In a recent study the rate of neurologic and ocular defects in fetuses born to ZIKV infected mothers were calculated to be ~7%^6^. Rarely, adults experience ZIKV-induced Guillain Barré syndrome^7^, but the congenital ZIka syndrome in infants that includes a spectrum of neurological defects, including microcephaly, is the most devastating and pressing aspect of ZIKV infection^8^. It is suggested that the risk of microcephaly is higher in the fetus when mothers are exposed to ZIKV during first trimester of pregnancy^3,6^. Some studies have reported the direct effects of ZIKV infection on neuronal tissue damage^9,10^. However, not all ZIKV infections during pregnancy appear to result in brain abnormality during embryonic development and the mechanisms that lead to microcephaly in some fetuses but not others are not yet understood.

Interestingly, ZIKV epidemic regions are often endemic for other Flavivirus infections, particularly DENV. Due to the structural similarities between ZIKV and DENV^11,12^, antibodies raised against one of the viruses are able to cross react with the others ^12,13^. It is thought that DENV antibodies can potentially lead to antibody-dependent enhanced replication (ADE) of ZIKV in infected patients^14^, which has been shown through *in vitro* assays^13^. The results for ADE in animal models of ZIKV infection have been less clear, with ADE apparent in immune-compromised mice but conflicting results for primates that showed no enhancement or very moderate enhancement of ZIKV infection in DENV-immune animals^15-17^. However, it has not been experimentally addressed whether DENV cross-reactive antibodies can influence vertical transmission of infection. Since maternal antibodies are also passed from mother to fetus during pregnancy in a mechanism dependent on FcRN^18,19^, which is expressed on the embryonic yolk sac, the fetal vascular endothelium, and syncytiotrophoblast cells^20,21^, this led us to hypothesize that DENV antibodies might enhance translocation of the virus to the developing fetus.

## Results

We characterized ZIKV infection in female C57BL/6 mice using the ZIKV strain H/PF/2013^22^ to determine if immune-competent mice would experience replicating ZIKV infection with this strain. After intra-peritoneal infection of mice with ZIKV, ZIKV RNA could be detected within 24h in the cells of the peritoneum (**Figure 1a**) and in lymphoid tissues, including the spleen (**Figure 1b**) and the mesenteric and iliac lymph nodes (**Figure 1c-d**). Detection of ZIKV persisted beyond 7 days at the site of infection and in lymphoid tissues of immune-competent mice (**Figure 1a-d**). Infectious ZIKV was also detected in the serum of mice by focus-forming assay (**Figure 1e**). We also observed splenic swelling, consistent with an inflammatory response (**Figure 1f**). However, in adult mice, we did not detect any ZIKV RNA in the brain (**Figure 1g**), which is consistent with our understanding that ZIKV causes only a mild febrile illness in the majority of adult human patients^4^. Our data showing that ZIKV RNA could be detected at the site of infection and in lymphoid tissues for at least 1 week by PCR supported that ZIKV infection persists long enough in immune-competent mice to allow assessment of its influence on fetal development during pregnancy.

**Figure 1:**
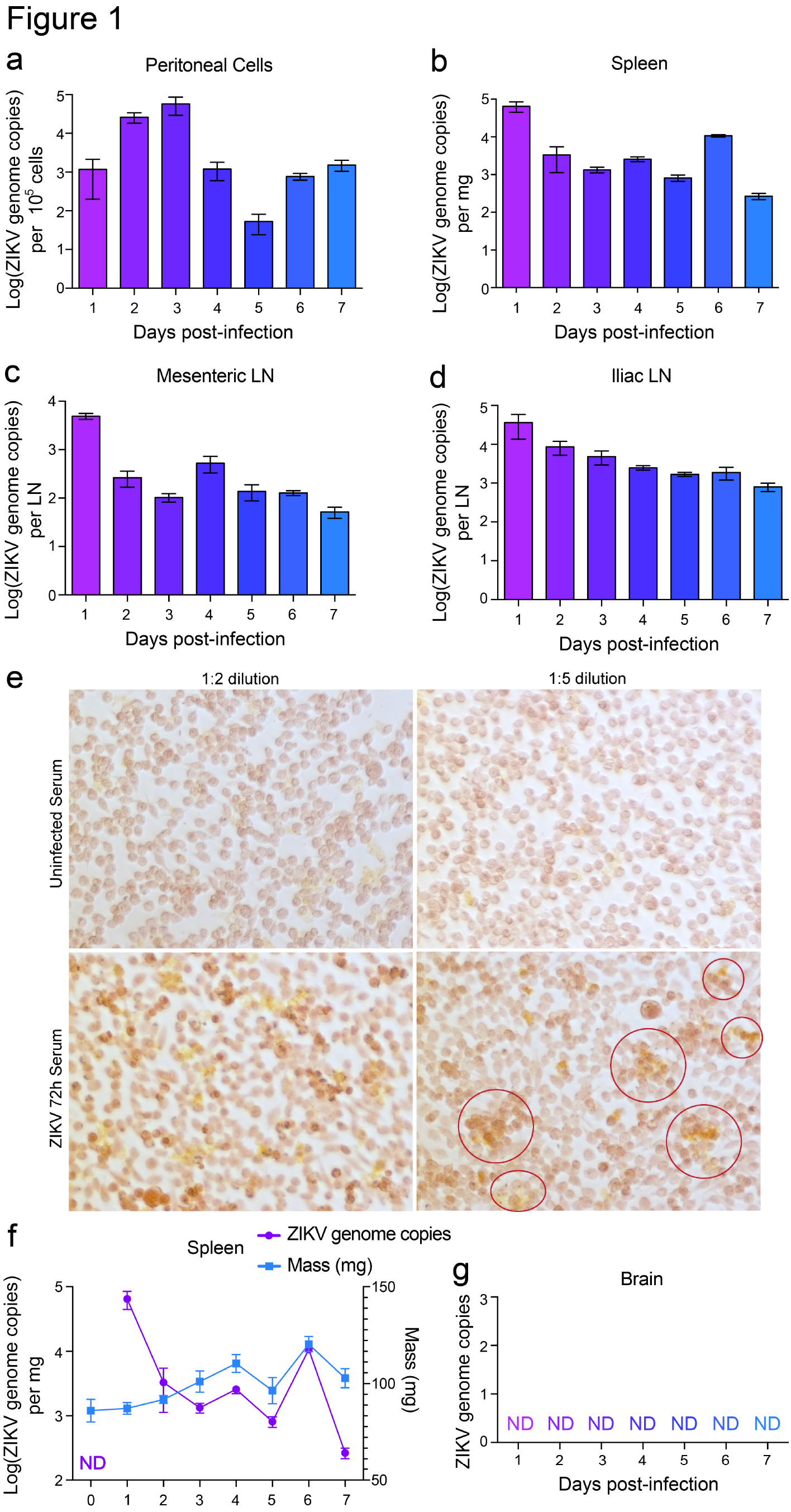
Sustained ZIKV infection in female immune-competent mice. ZIKV infection was quantified in peripheral tissues in (**a**) the cells isolated from the peritoneal cavity, (**b**) the spleen (**c**) the mesenteric LN and (**d**) iliac LNs, each day for 1w following infection of female mice with 1×10^6^ PFU of ZIKV, strain H/PF/2013. (**e**) Serum from uninfected or ZIKV-infected dams was harvested 72h post-infection. Many cells appear infected, staining above baseline levels of uninfected serum, with ZIKV-infected serum diluted 1:2. Individual foci of infection (red circles) surrounded by a few single infected cells can be observed at higher 1:5 dilution of ZIKV serum. (**f**) ZIKV genome copies in the spleen are plotted with the splenic mass. (**g**) ZIKV was not detected (ND) in the brain of any animals. N=5 female mice per group. Error bars represent the SEM.

To address the question of whether maternal DENV immunity could enhance embryonic neural complications, we infected mice of varying Flavivirus-immune status during pregnancy at embryonic day (E)-7 with ZIKV (**Figure 2a**). This day was chosen due to its equivalence to the first trimester of fetal development in humans and since it represents a time point when the mouse placenta has begun to develop, which starts around E5.5^23,24^. We also assumed that using a time point for inoculation where the placenta had not completely formed might facilitate vertical transmission of infection, while the fact that decidualization has begun by this time point^25^ would allow us to address the potential of fetal endothelial cells to promote antibody transport into the fetus. DENV induces a mild replicating infection in immunocompetent mice that induces Flavivirus cross-reactive antibodies^26,27^. Pregnant dams were either naïve (to both ZIKV and DENV), immune to DENV2 after clearing infection that preceded pregnancy by 3 weeks, or adoptively transferred monoclonal antibody 4G2, which was raised against DENV2 but is Flavivirus cross-reactive^28^. Adoptive transfer of 4G2 was used to identify the contributions of pre-existing cross-reactive antibodies alone (as a component of immunity) to ZIKV pathogenesis. DENV-immune mice were verified by ELISA to have DENV-specific antibodies that bind weakly to ZIKV (**Figure 2b**). At E18, near full-term, dams were euthanized and the fetal mice were examined. A gross analysis of the fetuses showed that mice that were previously infected with DENV had pups with stunted growth (**Figure 2c-d, Figure S1**). Quantification of the fetal mass showed a significant decrease in the mass of fetuses of naïve, ZIKV-infected dams, compared to those of naïve, uninfected controls (**Figure 2e**). This reduced body size was consistent with previous reports of ZIKV infection in mice^29,30^. However, infected DENV2-immune and 4G2-injected dams had fetuses that were even smaller than the naïve ZIKV-infected dams (**Figure 2c-e**). We examined the circumference of the fetal head, the primary measure used to define microcephaly in humans, and observed that DENV-naïve ZIKV-infected dams had fetuses with slightly but significantly smaller heads compared to healthy controls (**Figure 2f**). However, 4G2-injected and DENV2-immune dams that were infected with ZIKV had fetuses with head sizes that were substantially smaller than both naïve uninfected mice and DENV-naïve ZIKV-infected mice (**Figure 2f**). No significant differences in the range of phenotypes were observed litter-to-litter within experimental groups and fetal demise was not observed. Additional control experiments were also performed concurrently, showing that no abnormalities were observed after control injection of 4G2 prior to pregnancy without maternal ZIKV infection and isotype control antibodies did not influence fetal size or head circumference in ZIKV-infected dams (**Figure S2**). Furthermore, DENV-immune dams have normal pups without any reported fetal abnormalities^31,32^ and we did not observe any reduction in fetal size when 4G2-injected dams were given a challenge of DENV2 rather than ZIKV at E7 (**Figure S2**). Next, we looked at the frequency of microcephaly in these fetuses (defined as a head size in the 3rd percentile or less for normal fetuses) and noted that while 30% of fetuses from naïve mice infected at E7 with ZIKV could be assessed as having a phenotype consistent with microcephaly, >90% of the fetuses of DENV2-immune or 4G2-injected dams qualified as having microcephaly (**Figure 2g**). This supports that maternal immunity or Flavivirus cross-reactive antibodies can enhance the severity and incidence of microcephaly during ZIKV infection.

**Figure 2:**
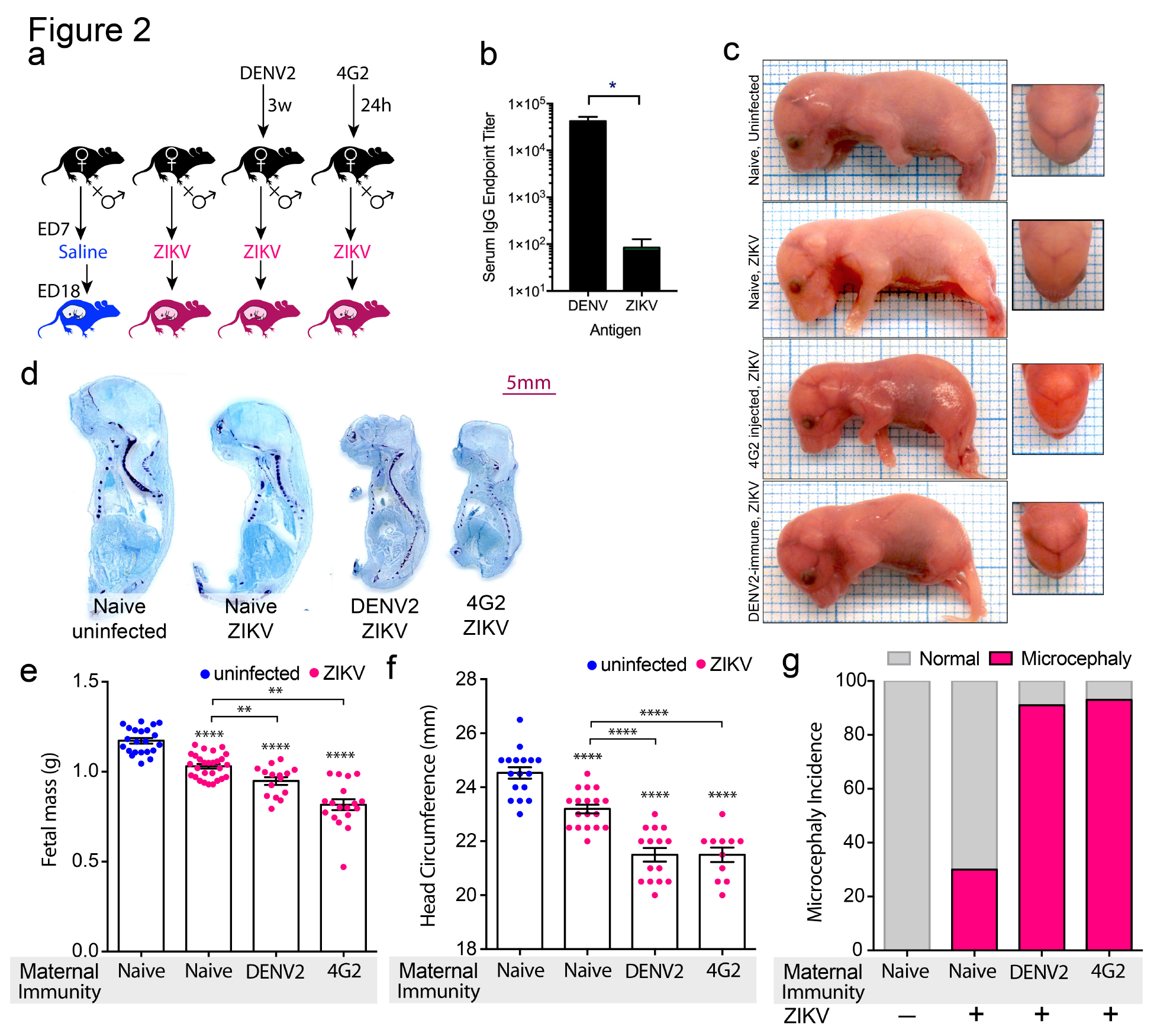
Maternal DENV immunity increases ZIKV-microcephaly in fetuses. (**a**) Schematic of the experimental time course showing that female naïve mice or mice that were infected with DENV2 three weeks prior or mice that were adoptively transferred the monoclonal antibody 4G2 (50μg, i.p.) were each crossed to male mice. Dams were infected on E7 after conception and the fetal development was assessed on E18. (**b**) ELISAs were performed against equivalent concentrations of DENV or ZIKV antigen using serum isolated 21d post-DENV infection. Serum from DENV-immune animals binds significantly more to DENV than to ZIKV. Graph represents the geometric mean titer 2-fold over naïve controls. Significance was determined by comparing endpoint dilutions by Student’s paired T-test; p=0.001. (**c**) Representative images of fetal mice on a 1-mm^2^ grid are provided to show that the DENV2-immune and 4G2-injected groups were visually smaller than controls. (**d**) Cross-sections of embryos stained with toluidine blue to reveal the reduced size of fetuses of DENV2-immune and 4G2-injected dams. Additional representative images are presented in **Figure S1**. (**e**) Mass and (**f**) head circumference of fetal mice on E18. (**g**) The incidence of mice having microcephaly, defined as a head size in the 3^rd^ percentile or less, calculated from the standard deviation of fetuses from naïve dams without ZIKV infection. For panels **e-f**, ^**^ p<0.01 and ^****^ p<0.0001, by 1-way ANOVA with Holm-Sidak’s multiple comparison test. Error bars represent the SEM. Variances do not differ significantly. For e, N=22 (naïve-uninfected), N=28 (naïve-ZIKV), N=15 (DENV2-ZIKV), N=18 (4G2-ZIKV). For **f**, N=17 (naïve-uninfected), N=18 (naïve-ZIKV), N=15 (DENV2-ZIKV), N=11 (4G2-ZIKV). Groups contain the combined data from all fetuses of 3 independent dams.

Due to the prominence of interferon-deficient models for studying Flaviviruses, we also investigated whether interferon (IFN)-deficient mice could be used to study ZIKV infection at time points developmentally analogous to the first trimester of human infection. However, even with a 2-log lower inoculating dose than we used for WT mice, the infection was too severe in both the naïve and DENV-immune dams. Most dams died prior to E18 (**Figure S3a**) and the surviving mouse had fetuses displaying early developmental arrest (**Figure S3b**). This supported that our immune competent model more clearly recapitulated the outcomes of human ZIKV maternal and fetal infection than the interferon-deficient mouse model system.

To further characterize the impact of ZIKV infection, with and without maternal antibodies on the development of the fetal brain in the WT mouse model, we examined the brains by histology (**Figure 3a, Figure S4**). The DENV2-immune group showed profound reduction in cortical thickness and loss of integrity of the expected cortical layers (**Figure 3a**). In particular, there appeared to be reductions in the size of the ventricular zone, intermediate zone and cortical plate compared to fetuses from naïve uninfected dams and naïve ZIKV-infected dams (**Figure 3a**). Quantification of cortical thickness from multiple litters showed that these reductions were significant and consistent for fetuses with both DENV2-immune and 4G2-injected dams compared to fetuses of naïve uninfected dams and naïve ZIKV-infected dams (**Figure 3b**). The cortical thickness was also moderately reduced in the fetuses of naïve ZIKV-infected dams compared to fetuses of healthy controls (**Figure 3b**), although the cortical layers were intact and discernable (**Figure 3a**). Since we had observed that the fetal mass was decreased in all ZIKV-infected groups compared to controls (**Figure 2d**), we also questioned whether the reduced cortical thickness was disproportionately small relative to the body mass or proportional to the body mass. Fetuses born to ZIKV exposed pregnant mothers have been described as displaying both proportional and disproportional microcephaly in humans^5^. Indeed, the ratio of the cortical thickness compared to the body mass for all groups showed that for ZIKV-infected mice, the cortex was disproportionately reduced compared to control animals (**Figure 3c, S5**), indicating a presentation of disproportionate microcephaly.

**Figure 3:**
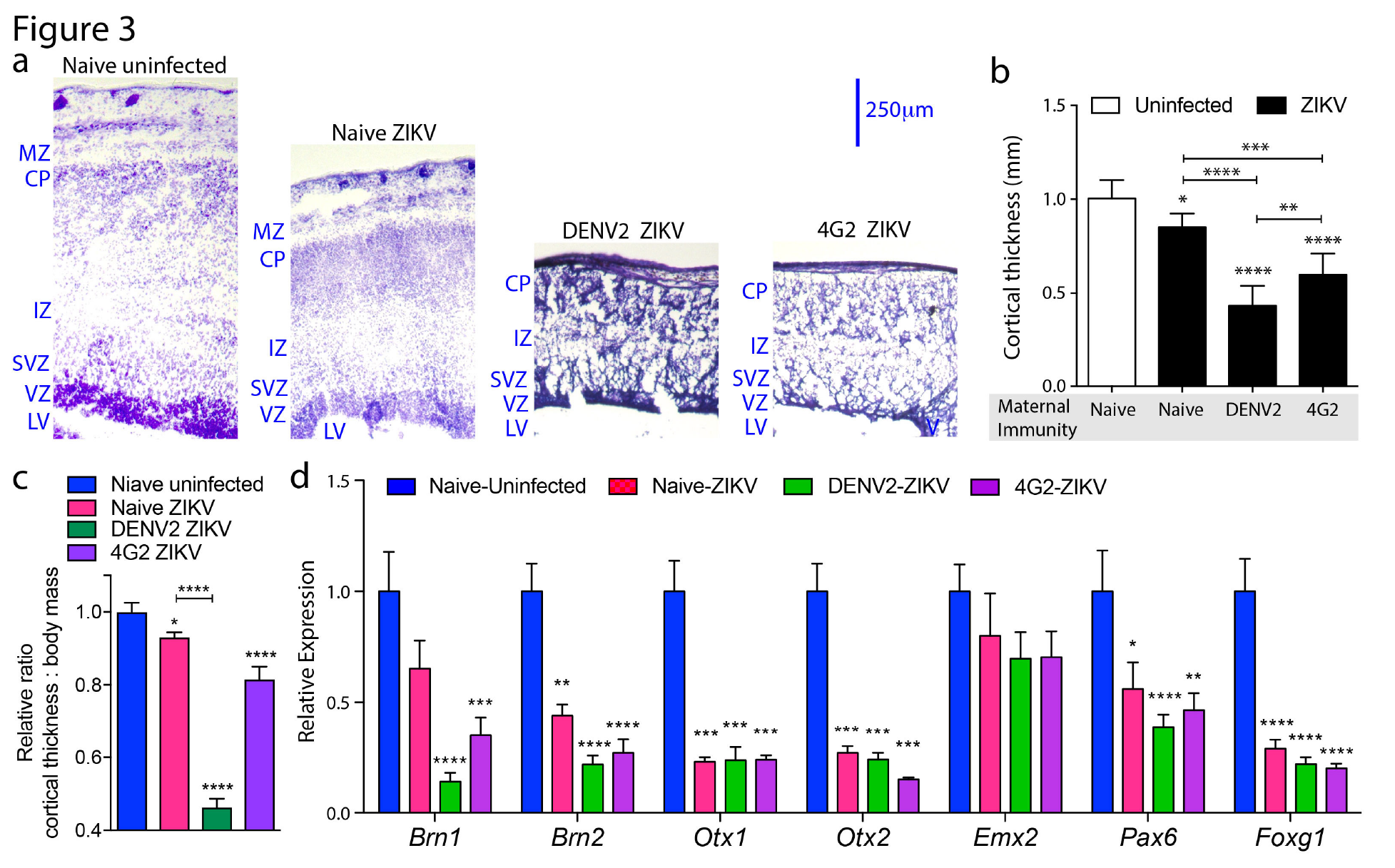
Impaired cortical development in ZIKV-fetuses is enhanced by maternal DENV immunity. (**a**) Images of embryonic brain sections showing reduced cortical thickness in ZIKV-infected embryos. Tissue sections were cut to 15μm thickness and nissl-stained using cresyl violet prior to imaging by light microscopy. Labels indicate the lateral ventricle (LV) and cortical regions: ventricular zone (VZ), sub-ventricular zone (SVZ), intermediate zone (IZ), cortical plate (CP) and marginal zone (MZ). Additional representative images are included in **Fig. S4**. (**b**) Quantification of cortical thickness from images. Error bars represent the SEM of average cortical thickness for individual fetuses obtained from 3 independent pregnancies; for naïve uninfected, n=5; naïve-ZIKV n=6; DENV2-ZIKV, n=7; 4G2-ZIKV, n=8. (**c**) Normalized ratio of cortical thickness to body mass for fetal mice. Additional controls are provided in **Fig. S5**. (**d**) Relative expression of transcriptional factors critical for neuronal growth in the fetal mouse brains (normalized to *Actin*) as determined by real time RT-PCR. Error bars represent the SEM and n=5 fetuses per group. For **b-c**, ^*^ indicates p<0.05, ^**^ p<0.01, ^***^ p<0.001 and ^****^ p<0.0001, as determined by 1-way ANOVA with Holm-Sidak’s multiple comparison test. Variances do not differ amongst groups.

The transcription factor Brain 1 (*Brn1*) is used as a marker of cortical development in mice^33^; thus, we measured levels of *Brn1* in the brains of E18 embryos from all groups. We observed that levels of *Brn1* mRNA were reduced in fetuses from DENV2-immune or 4G2-injected dams (**Figure 3d**), which was correlated with the impaired cortical thickness. We also measured levels of additional genes associated with early cortical neurogenesis, including fork-head box G1 (*Foxg1*), empty spiracles homologue 2 (*Emx2*) and Paired box 6 (*Pax6*). Expression of these genes allows differentiation and proliferation of ventricular zone progenitors as well as expansion of the sub-ventricular zone^34,35^. ZIKV infection resulted in reduced expression of both *Pax6* and *Foxg1*, suggesting early brain development was stunted due to ZIKV infection, although *Emx2* was not significantly influenced (**Figure 3d).** Brain 2 (*Brn2*) and orthodenticle homeobox protein family genes, *Otx1* and *Otx2*, along with *Brn1*, aid the differentiation and migration of neurons ^33,36^ as well as development of neuronal layers in the cortex and cerebellum ^37^. ZIKV infection also suppressed the expression of both *Brn2* and *Otx* family genes. For *Pax6*, *Brn1*, and *Brn2,* fetuses of DENV-immune dams showed greater deficits than those of naïve dams (**Figure 3d**). Suppressed levels of selected cortical markers were verified at the protein level by immunohistochemistry (**Figure S6**). Together these results show visually striking and quantifiable defects in the development of the cerebral cortex during ZIKV infection resulting in disproportionate microcephaly, supported by evidence that cerebral cortex-associated transcription factors are reduced in the fetuses of DENV2-immune dams.

Based on these findings and the potential of the antibody 4G2, which is DENV-directed but ZIKV cross-reactive, to cause an enhanced microcephaly-like phenotype in the fetuses of ZIKV-infected dams, we expected that antibodies might promote increased translocation of ZIKV into the fetus; therefore, we quantified ZIKV genome copies in the fetuses on E10, 3 days after the dams had been infected. This time was chosen to allow for replication of ZIKV in the mother and the potential of translocation into the fetus and because it is a time point at which placentation has already begun^23^. We first validated that true replication of ZIKV occurs in this model by comparing the infection levels in the dams and fetuses in mice injected with live ZIKV compared to UV-inactivated ZIKV. This demonstrated that input virus is only detected at very low levels in the dam’s spleen (**Figure 4a**), but not in the placenta (**Figure 4b**) or embryos (**Figure 4c**). The viral burden did not differ in pregnant mice compared to non-pregnant mice (**Figure S7**). Furthermore, the virus negative-strand, which is only produced during genome replication, was detectable by PCR in the dams’ spleens, the placentas, and also the embryos (**Figure 4d-e**), confirming ZIKV replication. These findings support that fetal ZIKV infection in this model relies on replicating ZIKV *in vivo*.

**Figure 4:**
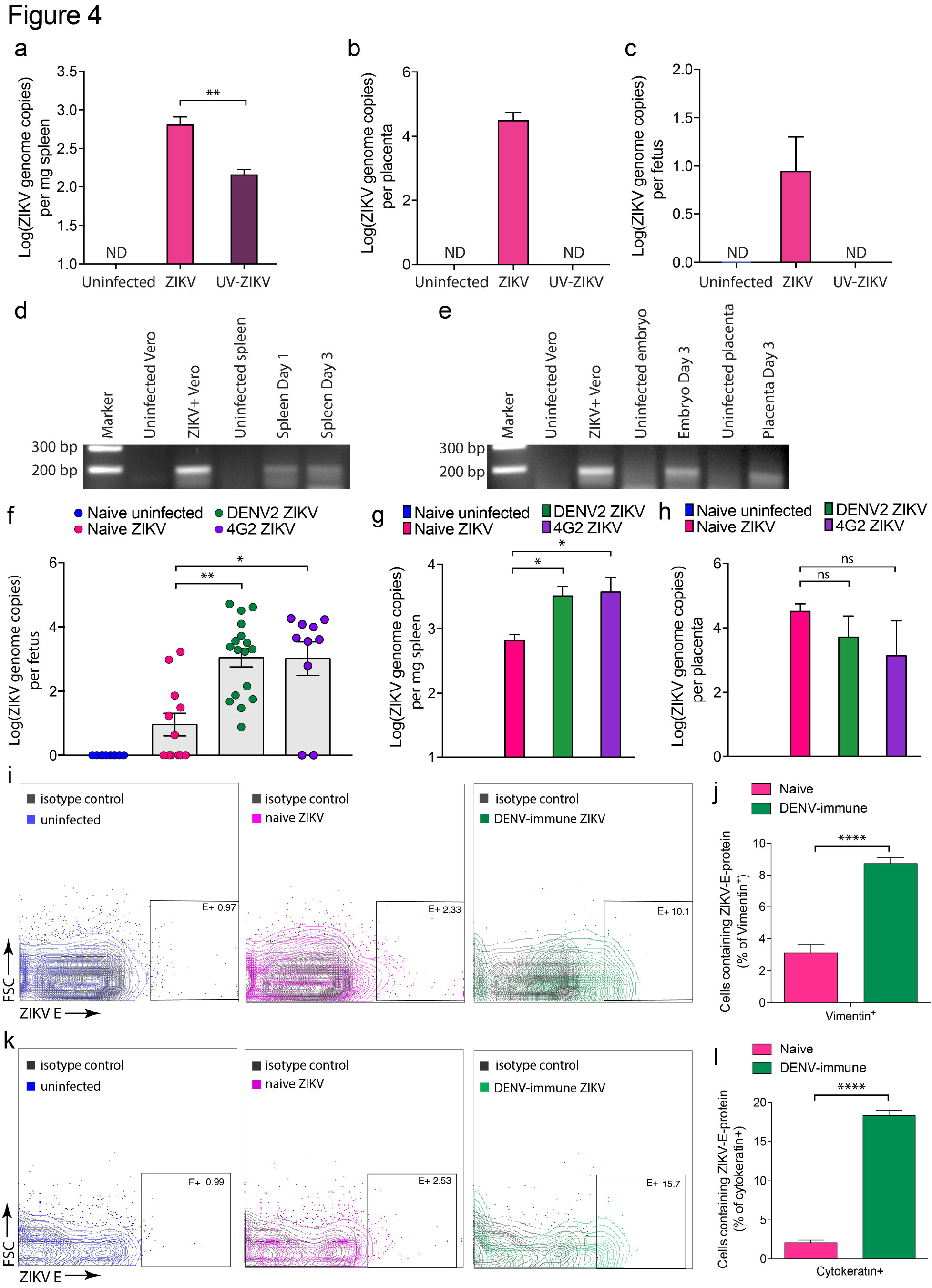
Enhanced ZIKV infection of fetuses in DENV-immune dams. (**a-c**) ZIKV genome copies were quantified by PCR after injection of UV-inactivation of virus, compared to injection of live virus in tissues including the (**a**) dam’s spleens (**b**) placentas or (**c**) fetuses on E10, 3d post-infection. UV-ZIKV could be detected only in the dam’s spleen, but at significantly lower genome copy numbers than live virus. PCR specific for the ZIKV negative-strand was performed using tissue from (**d**) the mother’s spleen days 1 and 3 post-infection and (**e**) from embryos or placentas on E10, 3d post-infection. For **d-e**, uninfected and ZIKV-infected vero cell lysates were used as negative and positive controls and uninfected tissues were used as negative controls. Expected band size is 188bp. The gels confirm active ZIKV replication in mother and fetal mice. (**f**) Real time RT-PCR was used to quantify the ZIKV genome copies in the mouse fetuses of DENV-naïve uninfected dams (n=10), from DENV-naïve ZIKV-infected dams (n=12), from DENV2-immune ZIKV-infected dams (n=17), and from 4G2-injected ZIKV-infected dams (n=10) derived from 2-3 independent experiments. Fetuses were harvested on E10, 3d after maternal infection or E18, 11d after maternal infection for RNA isolation. Fetuses from DENV2-immune and 4G2-injected dams showed significant increases in ZIKV compared to those of naïve dams by 1-way ANOVA with Holm-Sidak’s multiple comparison test (^*^p<0.01, ^**^p<0.001). (**g**) Quantification of ZIKV infection in the spleen of dams 3d after infection when the fetuses were harvested at E10 (n=3 per group). Error bars represent the SEM. (**h**) Quantification of ZIKV infection in the placentas at E10. (**i**) Representative flow cytometry plots showing intracellular staining for ZIKV E-protein in Vimentin^+^ fetal endothelial cells from the placenta at E10, following maternal ZIKV infection at E7. (**j**) E was detectable in a significantly larger proportion of Vimentin^+^ cells in the placentas from DENV-immune compared to naïve dams infected with ZIKV (n=5 per group). (**k**) Representative flow cytometry plots showing intracellular staining for ZIKV E-protein in cytokeratin^+^ synciotrophoblast cells from the placenta at E10, following maternal ZIKV infection at E7. (**l**) E was detectable in a significantly larger proportion of cytokeratin^+^ cells the placentas from DENV-immune compared to naïve dams infected with ZIKV (n=5 per group).

Our results examining the potential of maternal DENV immunity to influence fetal ZIKV infection showed that maternal antibodies to DENV or monoclonal antibody 4G2 each enhanced infection of the embryo compared to ZIKV infection of naïve dams (**Figure 4f**). ZIKV RNA was detectible in approximately 50% of the embryos of naïve, ZIKV-infected dams at E10 but 100% of the embryos of DENV2-immune and 80% of 4G2-injected dams (**Figure 4f**). Interestingly, by E18, although brain development was impaired (**Figure 3**), ZIKV RNA was undetectable in the brains of all fetuses by PCR; however, immunohistochemistry staining of E18 brain sections for ZIKV proteins NS2b and M showed positive staining, indicating that antigen persisted in the cortex (**Figure S8**). Enhanced detection of these antigens was observed in tissue sections from the fetuses of DENV-immune or 4G2-injected dams (**Figure S8**). DENV-immunity also enhanced ZIKV RNA in the dam’s spleen, consistent with other reports that DENV antibodies can cause ADE, we confirmed that this can occur *in vivo* in our model (**Figure 4g**). In contrast to the mother’s spleen, prior DENV immunity did not significantly enhance the viral burden in the placenta at E10 (**Figure 4h**). To validate this enhancement occurs *in vivo*, we performed flow cytometry staining for the ZIKV envelope protein (E) in single cell suspensions of placentas at E10, after E7 infection of dams. We observed that ZIKV E protein could be detected intracellularly in fetal endothelial cells, defined by vimentin staining (**Figure 4i-j**) and in synciotrophoblast cells, defined by the marker cytokeratin, increased over baseline staining of cells from uninfected control placentas (**Figure 4k-l**). Furthermore, the percentage of placental fetal endothelial cells and synciotrophoblasts that contained ZIKV E protein were significantly increased in the placentas of DENV-immune compared to naïve dams (**Figure 4i-l**). These results indicated that heterologous immunity enhances ZIKV infection in both the pregnant dam and the fetus and enhance detection of the ZIKV structural protein, E, in placental cells.

Since antibodies are translocated into the fetus with a mechanism dependent on FcRN, and we observed enhanced staining of E protein in the placentas of DENV-immune dams in two cell types that are known to express FcRN, we examined whether FcRN contributes to fetal ZIKV infection in DENV-immune mice. However, since there are conflicting statements in the literature regarding when FcRN expression begins in the fetus, we first verified that FcRN could be detected in fetal mice. Immunostaining of the mouse placenta at E10 (the time point when virus was detectable in mouse embryos by positive and negative-strand PCR, **Figure 4c,e,f**), revealed colocalization of FcRN with the fetal endothelial cell marker, vimentin (**Figure 5a**) and synciotrophoblast marker cytokeratin (**Figure 5b**). We next isolated mouse placentas from uninfected and ZIKV-infected mice at E8 and E10 to represent early time points, 24h and 72h post-maternal infection. We detected *FcRN* expression in infected and uninfected placentas at both E8 and E10 by PCR (**Figure 5c**). Interestingly, maternal ZIKV infection led to increased *FcRN* mRNA expression in E10 embryos, compared to controls (**Figure 5c**). This was validated by flow cytometry at E8 and E10, where nearly all fetal endothelial cells (vimentin^+^) and synciotrophoblasts (cytokeratin^+^) expressed FcRN (**Figures 5d-g**). Similarly to PCR detection, a slight but significant increase in FcRN protein was observed on the vimentin^+^ cells of placentas from ZIKV-infected dams on E8 compared to uninfected controls (**Figure 5d-e**). This induction of FcRN was more dramatic by E10 and apparent on both fetal endothelial cells and synciotrophoblasts (**Figure 5f-g**). These results demonstrate that FcRN is present in the placenta during and preceding the E10 time point when antibody-dependent enhanced detection of ZIKV is observed in the fetus (**Figure 4f**).

**Figure 5:**
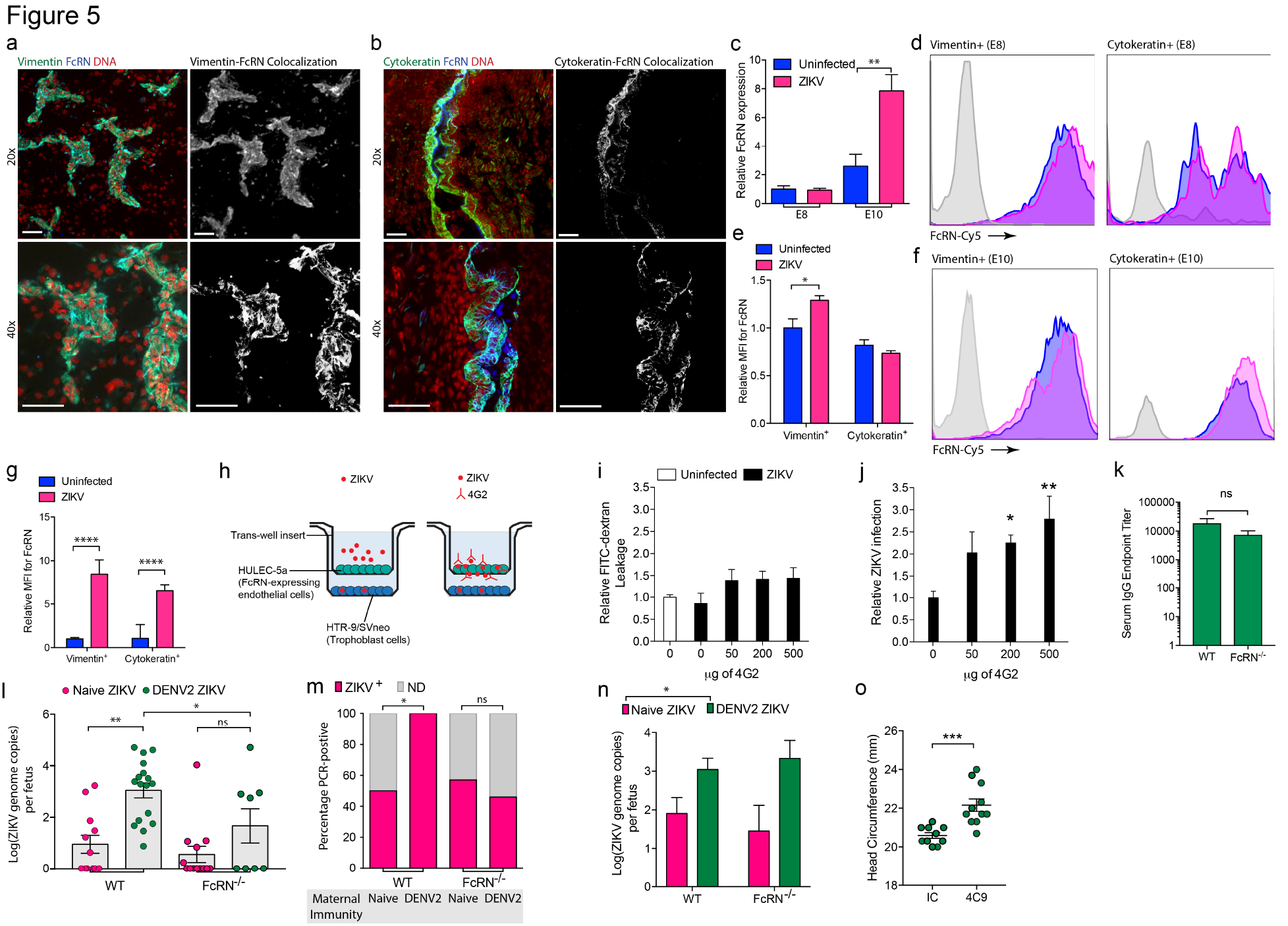
Maternal DENV immunity leads to FcRN-dependent enhancement of fetal ZIKV infection. Placenta tissue sections from E10 were stained for FcRN, DNA (DAPI), and either (**a**) the fetal endothelial cell marker vimentin or (**b**) the syncytiotrophoblast marker, cytokeratin and imaged by confocal microscopy. Both cytokeratin and vimentin co-localized with FcRN. Scale bars=50μm. Images are representative of 3 independent experiments. (**c**) FcRN mRNA expression in E8 and E10 mouse placentas. (**d**) Detection by flow cytometry of FcRN on cells from E8 placentas, gated on vimentin (left) or cytokeratin (right) from uninfected or ZIKV-infected dams, 24h post-infection at E7. (**e**) Average mean flouresence intensity (MFI) of FcRN on cells from E8 placentas, relative to levels on uninfected control endothelial cells (vimentin^+^); n=5 per group. (**f**) Detection by flow cytometry of FcRN on cells from E10 placentas, gated on vimentin (left) or cytokeratin (right) from uninfected or ZIKV-infected dams, 72h post-infection at E7. (**g**) Average MFI of FcRN on cells from E8 placentas, relative to levels on uninfected control endothelial cells (vimentin^+^); n=5 per group. (**h**) Schematic showing experimental set-up of trans-wells, where FcRN-expressing HULEC-5a cells were plated on trans-well inserts to form a tight monolayer, and trophoblast placenta cells were plated on the bottom chamber. ZIKV, or ZIKV+antibody 4G2 were added to the top chamber. (**i**) Permeability of the monolayers 24h after addition of ZIKV or ZIKV with various concentrations of antibodies was measured by quantitating FITC-dextran leakage. No differences in leakage were observed compared to uninfected controls. (**j**) Addition of antibody 4G2 significantly increased the amounts of infection in trophoblast target cells, in a dose-dependent manner, on the opposite side of the trans-well insert 24h post-exposure to ZIKV. (**k**) No significant difference was observed in serum endpoint titers four-fold over naive against DENV2 between WT and FcRN^-/-^ female mice 21d post-infection. P=0.32 by Student’s unpaired T-test, n=5 per group. (**l**) Quantification of ZIKV in DENV2-immune (n=17) or –naïve (n=10) WT and DENV2-immune (n=8) or –naïve FcRN^-/-^ (n=13) embryos on E10. (**m**) Proportions of PCR-positive embryos for each group in **c**. Significance was determined by Fisher’s exact test; ^*^p=0.01; ns for p=0.7064). (**n**) Comparison of ZIKV titers for ZIKV^+^ fetuses alone shows enhanced viral burden in the embryos of DENV2-immune dams for both WT and FcRN^-/-^ groups, determined by 1-way ANOVA with Holm-Sidak’s multiple compari son test; ^*^p=0.0043. For DENV-naïve WT, n=6; DENV2-immune WT, n=17; DENV-naïve FcRN^-/-^, n=5; DENV2-immune FcRN^-/-^, n=4, corresponding to the PCR-positive embryos from panel **l**. For **l-m**, all fetal measurements were derived from 3 independent dams, except for the uninfected controls, derived from 2 independent dams. (**o**) Blocking antibody against FcRN (4C9) or isotype control (IC) antibodies were injected therapeutically into DENV-immune pregnant mother mice prior to ZIKV infection on E7. Head circumference of the fetal mice was measured on E18. Head circumferences were significantly increased in 4C9-treated fetal mice compared to those given IC, p<0.0001.

To determine if ZIKV infection could be enhanced across a monolayer of FcRN-expressing endothelial cells in an antibody-dependent fashion, we using a trans-well system. Human endothelial cells were cultured above a monolayer of human trophoblast cells, and supernatants containing ZIKV or ZIKV and various concentrations of 4G2 antibody were applied to the top chamber (**Figure 5h**). After incubation for 24h, we assessed the permeability of the endothelial cell monolayers and observed no significant increase in permeability, measured by trans-endothelial resistance (**Figure 5i**). However, we observed an antibody-dose dependent increase in infection of the trophoblast target cells on the opposite side of the trans-well insert (**Figure 5j**). This indicates that antibodies enhance translocation of ZIKV across FcRN-expressing endothelial monolayers and that ZIKV remains viable and able to infect target cells following translocation.

We next aimed to determine if FcRN plays a functional role in enhancing ZIKV infection in fetal mice. First, we validated that FcRN^-/-^ mice were able to induce anti-DENV antibodies at similar titer to WT mice and observed no significant difference in the polyclonal anti-DENV response (**Figure 5k**). We then compared viral titers in fetuses of both naïve and DENV2-immune FcRN^-/-^ dams to wild-type controls. Overall, fetuses of DENV2-immune FcRN^-/-^ mice showed reduced ZIKV RNA at E10 compared to the fetuses of wild-type mice (**Figure 5l**). Furthermore, ZIKV RNA levels were not significantly higher in the fetuses of DENV2-immune dams compared to naïve dams (**Figure 5l**), in contrast to the significant enhancement of ZIKV infection observed in the context of DENV2 maternal immunity in wild-type animals (**Figure 5l**). In FcRN^-/-^ DENV2-immune dams, only 50% of the fetal mice showed ZIKV infection, compared to 100% of the fetuses of WT DENV2-immune dams (**Figure 5m**). This proportion was significantly lower than the proportion of infected fetuses in WT mice and not significantly different from ZIKV-infected naïve WT or naïve FcRN^-/-^ mice (**Figure 5m**) and suggests that FcRN-mediated translocation of immune complexes increases the likelihood of vertical transmission. Yet, antibodies may still have a role in enhancing titers in an FcRN-dependent manner, since DENV2-immune FcRN^-/-^ fetuses that were ZIKV^+^ showed higher viral titers than the DENV2-naïve FcRN^-/-^ fetuses that were ZIKV^+^ (**Figure 5n**). Similarly, the fetuses of wild-type dams that were DENV2-immune showed higher levels of ZIKV than naïve animals after ZIKV infection (**Figure 5n**), suggesting that DENV2-antibodies also result in enhanced levels of ZIKV if the fetus becomes infected by a mechanism independent of FcRN. To confirm the contributions of FcRN by an alternate mechanism, we also injected dams with a blocking antibody against FcRN to neutralize its function in the translocation of immune complexes across the placenta. We observed that fetal head circumference was increased with FcRN blockade, compared to isotype control-injection (**Figure 5o**). These results support that both a novel mechanism of FcRN-mediated immune complex translocation into the fetus and ADE can contribute to enhanced ZIKV infection in fetal mice.

## Discussion

Many viruses in the Flavivirus genus are neurotropic, but no others have been identified to cause microcephaly. Neurological complications occur only rarely with DENV^38^, the most closely related known human pathogen to ZIKV, and have entirely different clinical presentation in the context of Japanese encephalitis and West Nile viruses ^4^, involving encephalitis in adults and children. Some studies have reported neuronal tissue damage during ZIKV infection in humans and mice, involving infection of neural progenitor cells ^9,10^. Immuno-compromised mice develop severe fetal abnormalities ^30^, which are not necessarily consistent with the much lower rates of microcephaly that are observed in humans ^2^. Immune-competent mice infected with high doses of ZIKV during slightly later stages of pregnancy than our model also have fetal abnormalities, including reduced fetal size that corresponded with a reduced cortical thickness ^39^. Here, using a model of maternal infection at a time point of mouse fetal development corresponding to the first trimester of pregnancy in humans and a moderate ZIKV inoculating titer of a strain epidemiologically associated with microcephaly, we report that vertical transmission of ZIKV occurs. Furthermore, maternal antibodies enhance trans-placental infection of mouse fetuses and lead to exacerbated microcephaly. This involves reduced cortical thickness, substantial loss of certain cortical layers and impaired induction of transcriptional profiles key for brain development. Microcephaly in humans resulting from ZIKV infection has been described as both proportionate and disproportionate to the body mass ^5^. In our model, we observe disproportionate microcephaly relative to the reduction in overall fetal mass, which is enhanced by maternal DENV immunity. Infection was undetectable by PCR in E18 fetal brains while the signs of damage to the cortex and ZIKV antigens remained.

Head size alone did not indicate the full severity of reduced cortical thickness in mice, neither could this be interpreted from the pregnant dam’s viral burden, although ADE in the dams was evidenced by a higher ZIKV burden in DENV-immune compared to naïve pregnant mice. Our results also indicate that FcRN mediates a significant amount of the antibody-enhanced effects of disease since FcRN^-/-^ animals have reduced proportions of the embryonic mice infected with ZIKV *in utero*. This is likely due to the essential role of FcRN in mediating transcytosis of immune complexes across the placenta from mother to fetus ^40,41^ or across the syncytiotrophoblast cells prior to full placentation^23^. FcRN mediates transcytosis through recycling endosomes which are only weakly acidic and release of the antibody from FcRN occurs at neutral pH^42,43^; therefore, transcytosis of ZIKV-immune complexes is not expected to inactivate ZIKV since Flaviviruses are not inactivated until a pH <3.0^44^. Consistent with this, we observed enhanced infection of target trophoblasts in the presence of antibodies when ZIKV was exposed to the opposite side of a FcRN-expressing endothelial cell monolayer. We also confirmed the contributions of FcRN to maternal-fetal translocation of virus using an alternate method where FcRN was targeted with a monoclonal neutralizing antibody. In addition to supporting the role of FcRN in the antibody-enhanced phenotype in fetal mice, this experiment suggests the possibility of therapeutic targeting of FcRN to limit severe pathologies in the subset of mothers that are Flavivirus-immune. However, FcRN cannot account for all of the antibody-enhanced effects since embryos of DENV-immune FcRN^-/-^ dams that are infected have higher levels of ZIKV than naïve FcRN^-/-^ dams, indicating a contribution of ADE to enhanced fetal infection. In humans, maternal IgG is available in the fetus during the first trimester of human pregnancy ^45^ but the fetal concentrations of IgG are lower in the early (compared to late) stages of pregnancy ^45,46^. Thus, it will be important to determine if certain stages of human pregnancy are susceptible to antibody-enhanced translocation of virus and augmented microcephaly and whether the presence of Flavivirus-cross reactive antibody is a factor in extending the period of fetal susceptibility to ZIKV beyond the first trimester when most cases have been shown to occur ^5,47^. Additional studies are required to determine if DENV immunity is indeed a risk factor for microcephaly during maternal ZIKV infection in humans due to the inherent physiological and immune functional differences between humans and mice. A recent study examining this question found no association between the incidence of total abnormal pregnancy outcomes and positive tests for DENV-reactive IgG ^48^. However, since approximately 90% of the patients in that study were determined to be IgG positive, further studies are needed with a larger sample size of DENV IgG-negative patients to fully assess this question and to examine the effects of DENV-immunity on specific manifestations of ZIKV congenital syndrome, such as microcephaly. Addressing whether DENV immunity enhances severity of human ZIKV congenital syndrome is further complicated by the fact that ZIKV- and DENV-specific antibodies in serum cross-react to a large degree^49^, making it difficult to distinguish between immunity to ZIKV and DENV if a blood sample is not taken early enough after ZIKV infection to exclude the presence of DENV-cross-reactive ZIKV-antibodies. Genetic differences between Flaviviruses also play a strong role in the ability of antibodies raised against unique viruses to enhance versus neutralize a heterologous infection, as do the concentrations of virus-specific or cross-reactive antibodies. Thus, future studies are also needed to examine the potential of multiple strains and serotypes of DENV to enhance ZIKV-induced microcephaly since the incidence of microcephaly in humans is not high enough to suggest that 100% fetuses of DENV-immune mothers develop microcephaly. This study provides important considerations for microcephaly screening, raises novel therapeutic targets to limit severe disease, and informs our understanding of mechanisms that could influence the severity of ZIKV presentation.

## Methods

Female mice that were DENV-naïve or DENV-immune (3 weeks post-infection with 1×10^6^ PFU of Eden2 strain) were bred and infected with ZIKV (1×10^6^ PFU of H/PF/2013 strain) on E7. Detailed methods accompany the manuscript as a Supplementary Appendix.

## Acknowledgements

We thank the European Virus Archive for providing us with the ZIKV H/PF/2013 strain.

## References

1 Brasil, P. et al. Zika Virus Infection in Pregnant Women in Rio de Janeiro - Preliminary Report. N Engl J Med, doi:10.1056/NEJMoa1602412 (2016).

2 Johansson, M. A., Mier, Y. T.-R. L., Reefhuis, J., Gilboa, S. M. & Hills, S. L. Zika and the Risk of Microcephaly. N Engl J Med 375, 1–4, doi:10.1056/NEJMp1605367 (2016).

3 Cauchemez, S. et al. Association between Zika virus and microcephaly in French Polynesia, 2013-15: a retrospective study. Lancet 387, 2125–2132, doi:10.1016/S0140-6736(16)00651-6 (2016).

4 Musso, D. & Gubler, D. J. Zika Virus. Clin Microbiol Rev 29, 487–524, doi:10.1128/CMR.00072-15 (2016).

5 Brasil, P. et al. Zika Virus Infection in Pregnant Women in Rio de Janeiro. N Engl J Med 375, 2321–2334, doi:10.1056/NEJMoa1602412 (2016).

6 Hoen, B. et al. Pregnancy Outcomes after ZIKV Infection in French Territories in the Americas. N Engl J Med 378, 985–994, doi:10.1056/NEJMoa1709481 (2018).

7 Cao-Lormeau, V. M. et al. Guillain-Barre Syndrome outbreak associated with Zika virus infection in French Polynesia: a case-control study. Lancet 387, 1531–1539, doi:10.1016/S0140-6736(16)00562-6 (2016).

8 Mlakar, J. et al. Zika Virus Associated with Microcephaly. N Engl J Med 374, 951–958, doi:10.1056/NEJMoa1600651 (2016).

9 Wu, K. Y. et al. Vertical transmission of Zika virus targeting the radial glial cells affects cortex development of offspring mice. Cell Res 26, 645–654, doi:10.1038/cr.2016.58 (2016).

10 Culjat, M. et al. Clinical and Imaging Findings in an Infant With Zika Embryopathy. Clin Infect Dis, doi:10.1093/cid/ciw324 (2016).

11 Kostyuchenko, V. A. et al. Structure of the thermally stable Zika virus. Nature 533, 425–428, doi:10.1038/nature17994 (2016).

12 Barba-Spaeth, G. et al. Structural basis of potent Zika-dengue virus antibody cross-neutralization. Nature, doi:10.1038/nature18938 (2016).

13 Dejnirattisai, W. et al. Dengue virus sero-cross-reactivity drives antibody-dependent enhancement of infection with zika virus. Nat Immunol, doi:10.1038/ni.3515 (2016).

14 Halstead, S. B. Biologic Evidence Required for Zika Disease Enhancement by Dengue Antibodies. Emerg Infect Dis 23, 569–573, doi:10.3201/eid2304.161879 (2017).

15 Bardina, S. V. et al. Enhancement of Zika virus pathogenesis by preexisting antiflavivirus immunity. Science 356, 175–180, doi:10.1126/science.aal4365 (2017).

16 Pantoja, P. et al. Zika virus pathogenesis in rhesus macaques is unaffected by pre-existing immunity to dengue virus. Nat Commun 8, 15674, doi:10.1038/ncomms15674 (2017).

17 McCracken, M. K. et al. Impact of prior flavivirus immunity on Zika virus infection in rhesus macaques. PLoS Pathog 13, e1006487, doi:10.1371/journal.ppat.1006487 (2017).

18 Osborn, J. J., Dancis, J. & Rosenberg, B. V. Studies of the immunology of the newborn infant. III. Permeability of the placenta to maternal antibody during fetal life. Pediatrics 10, 450–456 (1952).

19 Israel, E. J., Patel, V. K., Taylor, S. F., Marshak-Rothstein, A. & Simister, N. E. Requirement for a beta 2-microglobulin-associated Fc receptor for acquisition of maternal IgG by fetal and neonatal mice. J Immunol 154, 6246–6251 (1995).

20 Simister, N. E., Story, C. M., Chen, H. L. & Hunt, J. S. An IgG-transporting Fc receptor expressed in the syncytiotrophoblast of human placenta. Eur J Immunol 26, 1527–1531, doi:10.1002/eji.1830260718 (1996).

21 Leach, J. L. et al. Isolation from human placenta of the IgG transporter, FcRn, and localization to the syncytiotrophoblast: implications for maternal-fetal antibody transport. J Immunol 157, 3317–3322 (1996).

22 Baronti, C. et al. Complete coding sequence of zika virus from a French polynesia outbreak in 2013. Genome Announc 2, doi:10.1128/genomeA.00500-14 (2014).

23 Watson, E. D. & Cross, J. C. Development of structures and transport functions in the mouse placenta. Physiology (Bethesda) 20, 180–193, doi:10.1152/physiol.00001.2005 (2005).

24 Sones, J. L. & Davisson, R. L. Preeclampsia, of mice and women. Physiol Genomics 48, 565–572, doi:10.1152/physiolgenomics.00125.2015 (2016).

25 Iguchi, T., Tani, N., Sato, T., Fukatsu, N. & Ohta, Y. Developmental changes in mouse placental cells from several stages of pregnancy in vivo and in vitro. Biol Reprod 48, 188–196 (1993).

26 Morrison, J. et al. Transcriptional Profiling Confirms the Therapeutic Effects of Mast Cell Stabilization in a Dengue Disease Model. J Virol 91, doi:10.1128/JVI.00617-17 (2017).

27 Saron, W. A. A. et al. Flavivirus serocomplex cross-reactive immunity is protective by activating heterologous memory CD4 T cells. Sci Adv 4, eaar4297, doi:10.1126/sciadv.aar4297 (2018).

28 Johnson, A. J., Martin, D. A., Karabatsos, N. & Roehrig, J. T. Detection of anti-arboviral immunoglobulin G by using a monoclonal antibody-based capture enzyme-linked immunosorbent assay. J Clin Microbiol 38, 1827–1831 (2000).

29 Li, C. et al. Zika Virus Disrupts Neural Progenitor Development and Leads to Microcephaly in Mice. Cell Stem Cell 19, 120–126, doi:10.1016/j.stem.2016.04.017 (2016).

30 Miner, J. J. et al. Zika Virus Infection during Pregnancy in Mice Causes Placental Damage and Fetal Demise. Cell 165, 1081–1091, doi:10.1016/j.cell.2016.05.008 (2016).

31 Martinez Gomez, J. M. et al. Maternal Antibody-Mediated Disease Enhancement in Type I Interferon-Deficient Mice Leads to Lethal Disease Associated with Liver Damage. PLoS Negl Trop Dis 10, e0004536, doi:10.1371/journal.pntd.0004536 (2016).

32 Ng, J. K. et al. First experimental in vivo model of enhanced dengue disease severity through maternally acquired heterotypic dengue antibodies. PLoS Pathog 10, e1004031, doi:10.1371/journal.ppat.1004031 (2014).

33 Sugitani, Y. et al. Brn-1 and Brn-2 share crucial roles in the production and positioning of mouse neocortical neurons. Genes Dev 16, 1760–1765, doi:10.1101/gad.978002 (2002).

34 Muzio, L. & Mallamaci, A. Foxg1 confines Cajal-Retzius neuronogenesis and hippocampal morphogenesis to the dorsomedial pallium. J Neurosci 25, 4435–4441, doi:10.1523/JNEUROSCI.4804-04.2005 (2005).

35 Muzio, L. et al. Conversion of cerebral cortex into basal ganglia in Emx2(-/-) Pax6(Sey/Sey) double-mutant mice. Nat Neurosci 5, 737–745, doi:10.1038/nn892 (2002).

36 McEvilly, R. J., de Diaz, M. O., Schonemann, M. D., Hooshmand, F. & Rosenfeld, M. G. Transcriptional regulation of cortical neuron migration by POU domain factors. Science 295, 1528–1532, doi:10.1126/science.1067132 (2002).

37 Frantz, G. D., Weimann, J. M., Levin, M. E. & McConnell, S. K. Otx1 and Otx2 define layers and regions in developing cerebral cortex and cerebellum. J Neurosci 14, 5725–5740 (1994).

38 Carod-Artal, F. J., Wichmann, O., Farrar, J. & Gascon, J. Neurological complications of dengue virus infection. Lancet Neurol 12, 906–919, doi:10.1016/S1474-4422(13)70150-9 (2013).

39 Cugola, F. R. et al. The Brazilian Zika virus strain causes birth defects in experimental models. Nature 534, 267–271, doi:10.1038/nature18296 (2016).

40 Roopenian, D. C. et al. The MHC class I-like IgG receptor controls perinatal IgG transport, IgG homeostasis, and fate of IgG-Fc-coupled drugs. J Immunol 170, 3528–3533 (2003).

41 Mathiesen, L. et al. Maternofetal transplacental transport of recombinant IgG antibodies lacking effector functions. Blood 122, 1174–1181, doi:10.1182/blood-2012-12-473843 (2013).

42 Huotari, J. & Helenius, A. Endosome maturation. EMBO J 30, 3481–3500, doi:10.1038/emboj.2011.286 (2011).

43 Walters, B. T. et al. Conformational Destabilization of Immunoglobulin G Increases the Low pH Binding Affinity with the Neonatal Fc Receptor. J Biol Chem 291, 1817–1825, doi:10.1074/jbc.M115.691568 (2016).

44 Jindadamrongwech, S., Thepparit, C. & Smith, D. R. Identification of GRP 78 (BiP) as a liver cell expressed receptor element for dengue virus serotype 2. Arch Virol 149, 915–927, doi:10.1007/s00705-003-0263-x (2004).

45 Jauniaux, E. et al. Materno-fetal immunoglobulin transfer and passive immunity during the first trimester of human pregnancy. Hum Reprod 10, 3297–3300 (1995).

46 Malek, A., Sager, R., Kuhn, P., Nicolaides, K. H. & Schneider, H. Evolution of maternofetal transport of immunoglobulins during human pregnancy. Am J Reprod Immunol 36, 248–255 (1996).

47 Li, R. Y. & Tsutsui, Y. Growth retardation and microcephaly induced in mice by placental infection with murine cytomegalovirus. Teratology 62, 79–85, doi:10.1002/1096-9926(200008)62:2<79::AID-TERA3>3.0.CO;2-S (2000).

48 Halai, U. A. et al. Maternal Zika Virus Disease Severity, Virus Load, Prior Dengue Antibodies, and Their Relationship to Birth Outcomes. Clin Infect Dis 65, 877–883, doi:10.1093/cid/cix472 (2017).

49 van Meer, M. P. A. et al. Re-evaluation of routine dengue virus serology in travelers in the era of Zika virus emergence. J Clin Virol 92, 25–31, doi:10.1016/j.jcv.2017.05.001 (2017).

